# Evolutionary trade-offs between antimicrobial resistance and virulence in *Pseudomonas aeruginosa*

**DOI:** 10.64898/2026.07.08.737271

**Authors:** Surajit Pal, Anton Mohn, Michael Habig, Barbara Pees, Hinrich Schulenburg

## Abstract

Antibiotics impose strong selection on bacteria, resulting in the emergence and spread of antimicrobial resistance. To date, the consequences of resistance evolution for other traits, especially virulence as a key life-history characteristic of high medical relevance remain poorly understood. A central limitation of existing work is reliance on clinical isolates with complex evolutionary histories, hindering causal inferences on resistance-virulence relationships. Such causal information is critical for our understanding of the distribution of resistance-associated evolutionary trade-offs, which may additionally guide optimization of treatment designs. The objectives of our study are to address these knowledge gaps using an experimental evolution framework with the human pathogen *Pseudomonas aeruginosa*. We quantified changes in resistance, life-history characteristics, *in-vivo* virulence with a *Caenorhabditis elegans* infection model, and whole genome sequences for bacterial clones, which had been independently evolved under three antibiotics with distinct cellular targets. We found that the resistance-virulence trade-off depended on the used antibiotic and was additionally driven by the underlying evolutionary path to resistance. Evolved resistance to the fluoroquinolone ciprofloxacin correlated positively with virulence, dependent on genetic changes in either the antibiotic target or efflux regulation. Conversely, evolved resistance to piperacillin/tazobactam, a β-lactam/β-lactamase inhibitor combination, was negatively correlated with virulence, contingent on the coincidental spread of resistance mutations and a large genomic deletion, containing numerous virulence genes. Lastly, evolved resistance to the aminoglycoside streptomycin led to only minor virulence changes. Overall, these antibiotic-specific evolutionary trajectories challenge the assumption of a universal resistance-virulence trade-off and demonstrate that antibiotic choice itself shapes pathogen virulence potential.

**Significance Statement:** Antimicrobial resistance and bacterial virulence are typically studied in isolation, yet they are shaped by the same evolutionary pressures and may directly compete for cellular resources. We addressed a fundamental but unresolved question: does evolution of resistance to an antibiotic make a pathogen more or less dangerous to a host? By characterizing experimentally evolved *Pseudomonas aeruginosa*, a leading cause of drug-resistant infections, we show that the answer depends on which antibiotic drove resistance. A positive resistance-virulence relationship occurred upon adaptation to a fluoroquinolone antibiotic, whereas it was negative upon β-lactam resistance, and unchanged upon aminoglycoside resistance evolution. These antibiotic-specific evolutionary trajectories challenge the assumption of a universal resistance-virulence trade-off and suggest that antibiotic choice shapes pathogen danger beyond drug susceptibility.

## Introduction

The current spread of antimicrobial resistance (AMR) represents a major threat to global health (1, 2). New treatment strategies are thus urgently required. An in-depth understanding of the relationship of AMR with other life history characteristics is of particular interest in this context, because it may help to identify promising targets as well as potential pitfalls of new treatment designs. For example, the evolution and expression of AMR often comes at a cost, creating a trade-off between survival under antibiotic exposure and competitive performance in antibiotic free environments or between resistance to one versus a second antibiotic (i.e., collateral sensitivity) (3–7). These trade-offs may offer opportunities for treatment optimization by inclusion of antibiotic-free time periods or through the combination of antibiotics that express collateral sensitivities towards each other (8, 9). Another medically highly relevant, yet generally neglected trade-off concerns the relationship between AMR and virulence. We here focus on virulence defined as pathogen-induced host mortality rate. This ability of an infecting pathogen to cause host mortality can be mediated through the production of toxins, and/or the colonization of the host as well as the subsequent proliferation of the pathogen within the host. To date, the large majority of studies on AMR do not specifically characterize virulence. Standard diagnostics include tests on AMR but not virulence. Laboratory-based studies on AMR of human pathogens usually do not assess virulence, because suitable animal models for virulence tests are not always established and/or virulence cannot always be easily predicted through host-free molecular and genomic characterizations. The few available studies suggest that AMR evolution can be accompanied by changes in virulence (10–16). Notably, several molecular pathways underlying AMR intersect with quorum sensing (QS)-regulated systems that regulate expression of virulence factors. For instance, efflux pumps can export both antibiotics and QS signaling molecules, while mutations affecting antibiotic targets or cell-envelope structure can induce widespread regulatory rewiring, altering stress responses and toxin production (17, 18). These overlaps imply that AMR evolution may have pleiotropic effects on pathogenicity, yet the direction of the relationship is usually unclear. In fact, in the available studies, AMR evolution was found to either attenuate, maintain, or even enhance virulence (10–16). As a result, predicting how AMR evolution shapes virulence remains a major unresolved puzzle in microbial evolution and infectious-disease biology.

A key challenge of most existing studies is their focus on clinical isolates with complex and often unresolved evolutionary histories. Because such strains accumulate numerous mutations under heterogeneous selective pressures, it is difficult to disentangle the direct effects of evolved AMR from unrelated host-adaptation processes. Moreover, virulence is often inferred from isolated *in-vitro* traits or via identifying specific genetic signatures, which capture only limited aspects of the infection process and commonly fail to predict outcomes *in-vivo*. Experimental evolution provides a powerful framework to overcome these challenges by enabling adaptation under controlled conditions with known evolutionary trajectories. This approach has been used repeatedly to reconstruct the evolution and molecular basis of resistance towards distinct antibiotic treatment designs with a variety of pathogens (3, 19, 20). In principle, virulence could be assessed for the experimentally evolved bacteria with the help of informative infection models, thereby enabling the inference of potential causal links between AMR evolution and virulence outcomes. To date, however, such inferences have not been used to assess how precisely the evolution of AMR causes a change in virulence.

The objectives of the current study are to address this current knowledge gap. We use a controlled experimental approach to dissect the possible causal links between AMR evolution and virulence in a highly problematic human pathogen, the Gram-negative bacterium *Pseudomonas aeruginosa*, as a model. This opportunistic one-health pathogen infects diverse hosts, including humans, livestock, and plants. It employs distinct infection strategies ranging from toxin-mediated acute virulence to persistent, biofilm-associated chronic infections (21). This pathogen represents a particularly critical threat for humans, because it often expresses multidrug resistance, while also showing a high potential for rapid adaptation (21, 22). Moreover, comprehensive knowledge is available on the genetics and molecular basis of both AMR and virulence that can be used as reference for the experimental work. For the purposes of our study, we leverage highly resistant clones previously generated through parallel experimental evolution of *P. aeruginosa* exposed to three clinically relevant antibiotics with distinct modes of action: ciprofloxacin (CIP), piperacillin/tazobactam (PIT), and streptomycin (STR) (23). We performed an in-depth characterization of independently evolved clones per antibiotic, combining AMR profiling, whole genome sequence analysis, measurements of growth as a proxy to fitness, *in-vitro* virulence factor production, and also the analysis of *in-vivo* virulence with the help of a *Caenorhabditis elegans* infection model. The nematode *C. elegans* has previously been established as a highly informative model to assess both acute and chronic infection with *P. aeruginosa* that recaptures most virulence mechanisms of relevance for human infections (24, 25). Our integrative analysis enables a systematic dissection of how distinct resistance mechanisms influence virulence within a controlled genetic background. Our results challenge the assumption of a universal resistance–virulence trade-off. Instead, we identified antibiotic-specific evolutionary trajectories that differentially reshape virulence.

## Results

### Evolved AMR affects resistance to other antibiotics and general growth characteristics

Our analysis is focused on nine to ten clones which we isolated from independently evolved populations from a previous evolution experiment (23) that were exposed to either CIP (10 clones from independently evolved replicate populations), PIT (10 clones), and STR (9 clones). We determined the minimum inhibitory concentration (MIC) for the three antibiotics and confirmed that the clones expressed high resistance to the respective antibiotic which they experienced during experimental evolution (Fig. 1A, Supplement Statistical Results). In addition, six out of ten CIP-evolved clones showed significant collateral sensitivity (i.e., higher susceptibility than expressed by the ancestor) to the STR aminoglycoside, consistent with our previous study and also other reports (23, 26–28). Variable responses were expressed by these clones towards PIT. Evolved PIT resistance led to significant cross-resistance to the other two antibiotics for most clones, and only four significant cases of collateral sensitivity to STR. Evolved STR resistance led to either non-significant resistance changes towards PIT, and almost always significant cross-resistance towards CIP. Together, these data demonstrate that collateral outcomes are drug-pair dependent and clone specific, while the collateral sensitivity trade-off is most consistently expressed upon evolution of CIP resistance and a few cases of PIT resistance.

**Figure 1.**
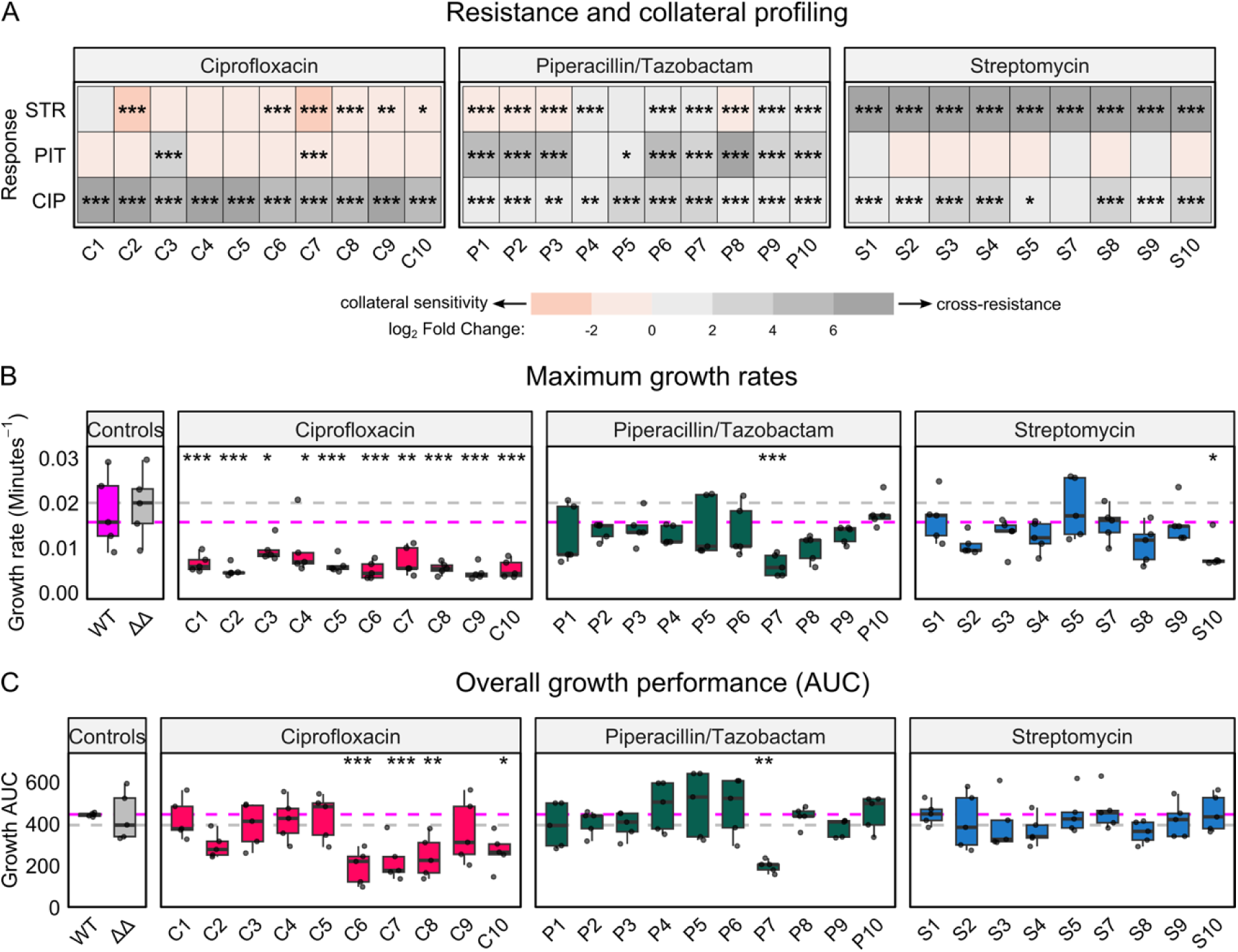
Resistance and collateral profiling of evolved *P. aeruginosa* clones and their growth performance in an acute-killing environment. (A) Collateral resistance and sensitivity profiles of antibiotic-evolved *P. aeruginosa* PA14 clones. The heatmap summarizes the collateral responses of individual clones evolved under CIP, PIT, or STR, when challenged with these same antibiotics as mean of 3 biological replicates. Colors represent the relative change in resistance compared to the ancestral wild-type (WT) PA14 strain, allowing classification of collateral sensitivity (red), cross-resistance (grey), or neutral responses (white) for each antibiotic combination. Statistical analysis was performed using Dunnett’s test for multiple comparisons, where each evolved strain’s MIC fold change and collateral responses were calculated relative to the ancestral WT. (B-C) Growth performance of evolved clones in the medium used for the *C. elegans* acute-killing assay, serving as a proxy for bacterial fitness (n = 5). Quantified parameters include (B) maximum growth rate (μ) and (C) cumulative growth performance (area under the curve, AUC). The magenta dashed line represents the WT PA14 reference and the grey dashed line indicates the *ΔlasR ΔrhlR* null mutant (ΔΔ) as negative control. Statistical comparisons were performed using Dunnett’s multiple comparison test against the WT PA14 strain. Significance thresholds are indicated as \**p* < 0.05, \*\**p* < 0.01, and \*\*\**p* < 0.001 (see also Supplement Statistical Results and additional results on growth performance in Figs. S1, S2).

To further quantify physiological costs of evolved AMR, we measured growth characteristics of the clones in antibiotic-free media, identical to the later used media for the acute and chronic *C. elegans-Pseudomonas* infection assays. Growth was recorded by measuring OD_600_ for 24 h in a plate reader, and dynamics were summarized as maximum growth rate, lag time, carrying capacity, and area under the curve (AUC) of the growth performance over time. We found that CIP-evolved clones showed consistent and significant reductions in maximum growth rates, prolonged lag phases and lower overall growth performance relative to the ancestral WT. These variations are seen in both media, although to a larger extent in the acute killing medium (Figs. 1B, 1C, S1, S2; see also Supplement Statistical Results). The observed reductions indicate a reproducible growth cost associated with CIP resistance, consistent with several previous studies (23, 29, 30). PIT-selected clones largely retained near-ancestral growth dynamics, although one PIT clone (clone P7) showed markedly slower growth and longer lag in both acute and chronic media, while half of the PIT isolates exhibited reduced AUC specifically in the chronic-assay medium (Figs. 1B, 1C, S1, S2; see also Supplement Statistical Results). STR-evolved clones were, for the most part, indistinguishable from the ancestor across all measured parameters, with the exception of clone S10, which produced a significantly slower growth rate in the acute medium, indicating minimal pleiotropic cost under our assay conditions (Figs. 1B, 1C, S1, S2; see also Supplement Statistical Results). Together, these data show that the fitness consequences of evolved AMR vary by antibiotic class and media composition: CIP resistance imposed the largest and most consistent growth costs, whereas PIT and STR resistance produced smaller or more heterogeneous effects that were often condition-dependent.

**Figure 2.**
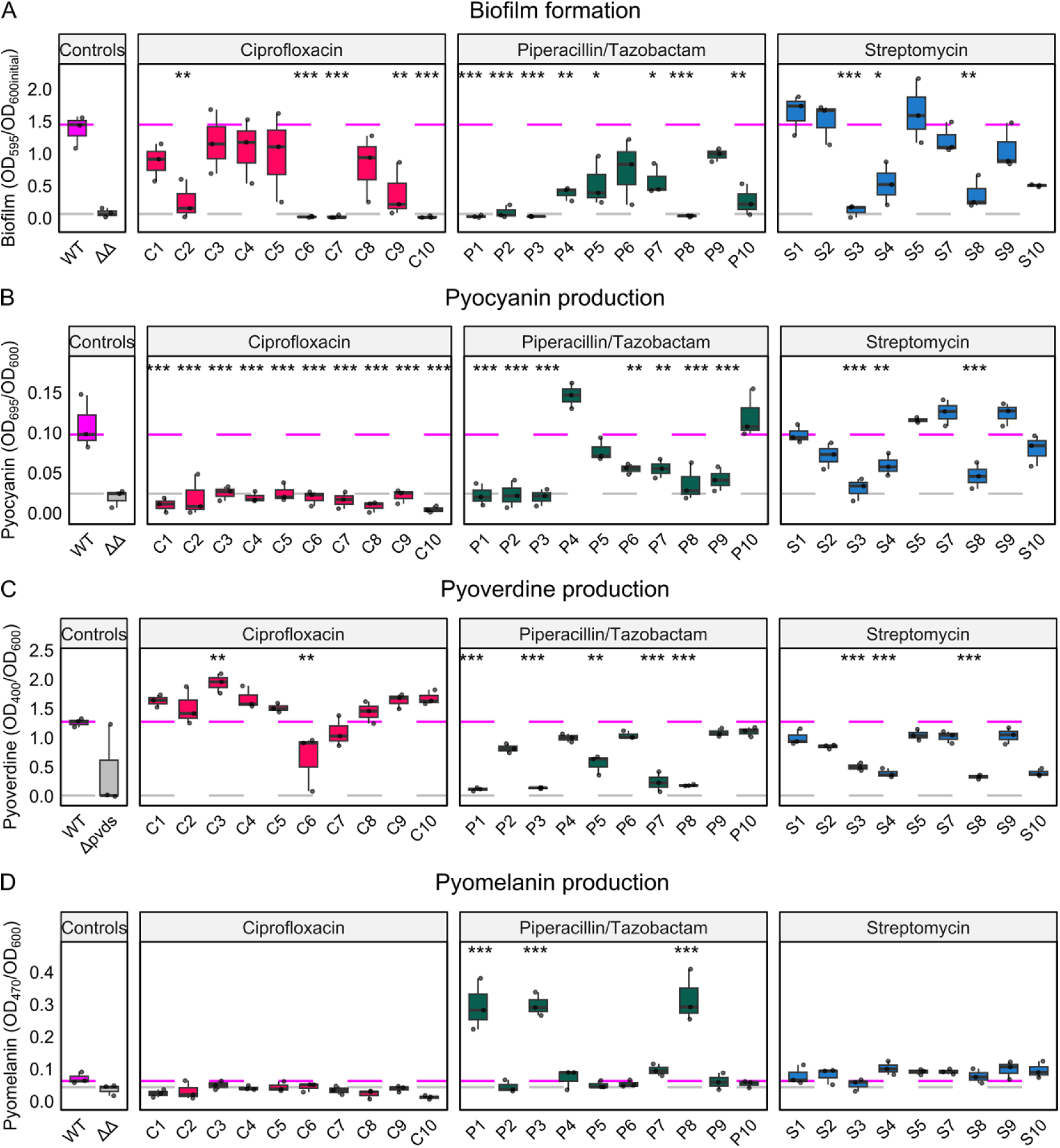
*In-vitro* quantification of virulence factor and pyomelanin production across antibiotic-evolved *Pseudomonas aeruginosa* clones. (A) Biofilm formation (n = 3), measured as the absorbance of crystal violet-stained adherent biomass at 595 nm and normalized to initial culture density (overnight cultures diluted to OD_600_ = 0.1). (B) Pyocyanin production (n = 3), quantified from cell-free supernatants at 695 nm and normalized to culture density (OD_600_). (C) Pyoverdine production (n = 3), assessed by regrowth of overnight cultures in iron-limited M9 medium and quantified at 400 nm. (D) Pyomelanin production (n = 3), quantified from cell-free supernatants at 405 nm and normalized to culture density (OD_600_). In all panels, the magenta dashed line denotes the median value of the wild-type PA14 strain (WT), the grey dashed line the median of (A–C) the *ΔlasR ΔrhlR* (ΔΔ) or (D) the *ΔpvdS* mutant as negative controls. Statistical comparisons were performed using Dunnett’s test against the WT. Significance thresholds are indicated as \**p* < 0.05, \*\**p* < 0.01, and \*\*\**p* < 0.001 (see also Supplement Statistical Results).

### Evolved AMR alters the expression of known *Pseudomonas* virulence traits

To test whether AMR evolution perturbs QS–dependent virulence traits, we quantified *in-vitro* biofilm formation and pyocyanin production. While biofilms can enable pathogen persistence within a host, pyocyanin is a redox-active toxin associated with acute and chronic virulence. We found significantly reduced biofilm production in half of the CIP-evolved clones, eight out of the ten PIT-resistant clones, and three STR-resistant clones (Fig. 2A; see Supplement Statistical Results). The changes in pyocyanin production were even more pronounced. All CIP-evolved clones lost their ability for pyocyanin production entirely, indicating a near-complete disruption of this QS-regulated toxin (Fig. 2B; Supplement Statistical Results). Seven out of the ten PIT clones similarly produced significantly less pyocyanin (Fig. 2B), while STR clones retained ancestral pyocyanin levels except for three significant reductions in the clones S3, S4, and S8, which also already showed significant reductions in biofilm formation (Fig. 2A, 2B). Because both biofilm formation and pyocyanin production are controlled by the Las/Rhl/PQS quorum-sensing circuits in *Pseudomonas*, these traits commonly covary (31). In our dataset, STR-evolved clones and many PIT-evolved clones retained this link. Conversely, CIP-evolved clones showed a clear decoupling of this relationship: pyocyanin secretion was abolished or strongly reduced while biofilm phenotypes were heterogeneous (Fig. 2A, 2B).

To complement the QS-regulated traits, we measured pyoverdine, the primary high-affinity siderophore of *P. aeruginosa*, which contributes to virulence by affecting bacterial iron acquisition and thereby host colonization (32). Pyoverdine production varied with selection history (Fig. 2C): half of the PIT clones and three STR clones produced significantly less pyoverdine than the ancestor, whereas CIP clones showed bidirectional responses, including a significant increase in pyoverdine production for clone C3, and a significant decrease for C6 (Fig. 2C; Supplement Statistical Results). These results indicate a more complex consequence of AMR evolution on pyoverdine production and iron scavenging. Interestingly, the three indicated STR clones with significantly decreased pyoverdine production are the same that also express a significant decrease in their ability to form biofilms and produce pyocyanin, suggesting a mechanistic link in the expression of these traits upon the evolution of resistance against STR only.

As an additional virulence trait, we also quantified the production of pyomelanin, a stress-associated pigment linked to oxidative protection and persistence (33). Only three clones showed a significant change in pyomelanin production, all with evolved PIT resistance (clones P1, P3 and P8; Fig. 2D; Supplement Statistical Results). These same clones also produced very high PIT resistance (MIC > 32×), while showing significant reductions in biofilm formation, pyocyanin and pyoverdine production. The combined pattern (i.e., no pyoverdine and pyocyanin with increased pyomelanin production) suggests some kind of mechanistic link between AMR and these virulence-related traits.

### Antibiotic-specific AMR effects on acute and chronic *C. elegans* infection characteristics

To test whether the *in-vitro* changes in virulence traits translate into altered *in-vivo* virulence, we quantified host mortality during both acute and chronic infection, using an established *C. elegans–P. aeruginosa* infection model (24, 25). The acute assay captures rapid, toxin-mediated killing over hours, whereas the chronic assay measures slower, persistence-dependent mortality. To assess possible biases due to bacterial densities, we ran acute infections under two inoculum regimes (high inoculum, using undiluted overnight cultures; and low inoculum, using cultures diluted to OD_600_ = 0.1). We confirmed by counting colony-forming units (CFUs) that the low-OD treatment produced substantially lower bacterial densities on assay plates (Fig. S3B). Nevertheless, both low and high inoculum treatments produced comparable virulence patterns, including comparable rank-order in virulence measures (Fig. 3A, Figs. S3A, S3C). Because of these identical outcomes, we focused on the high-inoculum protocol for both acute and chronic infection assays (Fig 3A, 3B; see also Supplement Statistical Results).

**Figure 3.**
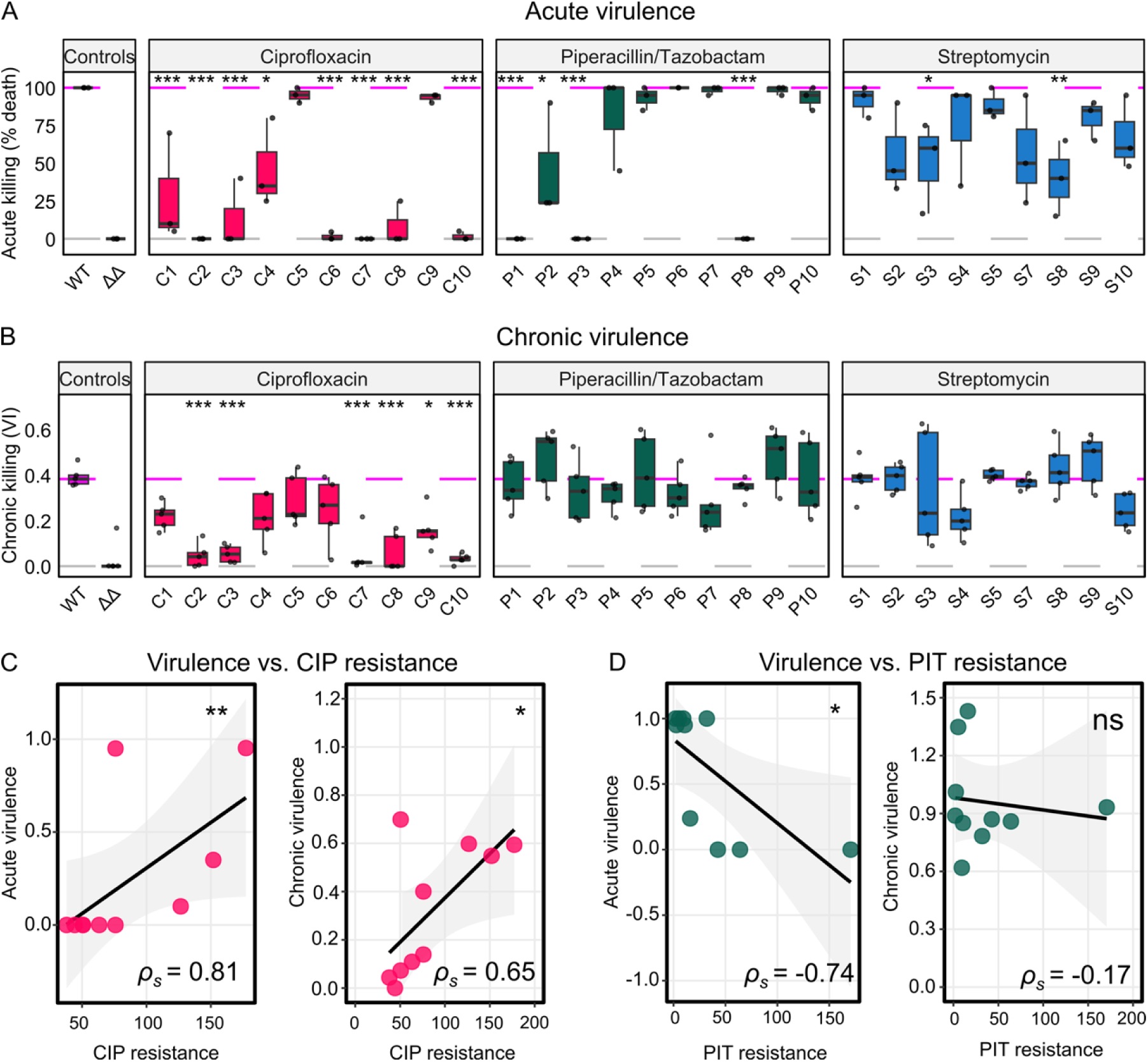
*In-vivo* virulence of antibiotic-evolved *P. aeruginosa* clones and its correlation with antibiotic resistance. (A) Results of the acute-killing assay that is driven by toxin-mediated virulence and host mortality occurs within hours of exposure; virulence is shown as the percentage of worms dead after 6 h post infection (n = 3). (B) Results of the chronic-killing assay, performed on low-osmolarity medium, for which virulence mainly results from persistent colonization and exploitation of host resources. In panels (A) and (B), the magenta dashed line denotes the wild-type PA14 strain (WT), and the grey dashed line the *ΔlasR ΔrhlR* double quorum-sensing mutant (ΔΔ), which serves as an avirulent negative control. Statistical comparisons were performed using Dunnett’s test against the WT. (C) Correlations between *in-vitro* antibiotic resistance (fold change in MIC) and *in-vivo* virulence for clones evolved under CIP (acute virulence is shown in the left panel, chronic virulence in the right panel). Spearman rank correlation; n = 10. (D) Correlations between *in-vitro* antibiotic resistance (fold change in MIC) and *in-vivo* virulence for clones evolved under PIT (acute virulence is shown in the left panel, chronic virulence in the right panel). Spearman rank correlation; n = 10. Significance thresholds are indicated as \**p* < 0.05, \*\**p* < 0.01, and \*\*\**p* < 0.001 (see detailed statistical results in the Supplement Statistical Results file). Additional results on chronic virulence are shown in Fig. S3.

Virulence varied substantially by resistance background. Although most CIP-evolved clones showed reduced acute killing consistent with loss of QS outputs (Fig. 3A; Fig. 2B), a subset (notably C5, C9) retained wild-type killing rates. PIT-evolved clones were highly heterogeneous: Three isolates (P1, P3, P8) showed a highly significant loss of acute virulence, while almost all others maintained ancestral toxicity. STR-evolved clones largely retained acute virulence levels, with two exceptions (S3, S8) that showed significantly attenuated killing (Fig. 3A).

Chronic outcomes diverged from acute patterns (Fig. 3B). For both PIT and STR clones, we did not find any significant difference to ancestral virulence under chronic infection. A significant reduction in chronic virulence was only observed for six CIP-evolved clones (Fig. 3B). Five of these also showed significantly lower acute killing. These results show that toxin-mediated and chronic virulence are likely mediated by independent mechanisms and only overlap in a few cases, dependent on AMR history.

To further examine the latter point, we compared the evolved AMR patterns, represented as MIC fold change relative to the ancestral strain, with the inferred *in-vivo* virulence measures for both infection modes. STR-evolved clones were excluded from this analysis because all isolates exceeded the upper quantification limit of the MIC assay (MIC > 1024 g/L), preventing meaningful discrimination of resistance levels. CIP-evolved clones exhibited a significant positive correlation between resistance and virulence across both acute and chronic infection contexts (Fig. 3C, S3C; Supplement Statistical Results), indicating that highly resistant isolates tended to retain pathogenic potential. In contrast, PIT-evolved clones showed a significant negative correlation between resistance and virulence in the acute infection model, consistent with a resistance–virulence trade-off upon PIT selection (Fig. 3D; Supplement Statistical Results). No significant correlation was detected in the chronic infection model (Fig. 3D), suggesting that persistent infection in PIT-selected backgrounds does not depend on resistance level. Overall, our results revealed two opposite types of the AMR–virulence relationship that was contingent on the AMR evolution background.

### Distinct genetic changes underlie AMR evolution

To identify the genetic changes underlying AMR evolution and resulting effects on growth and virulence, we sequenced the genomes of all 29 evolved clones together with the ancestral *P. aeruginosa* PA14 strain. Mutations across CIP-, PIT-, and STR-selected backgrounds are summarized as a gene-level heatmap (Fig. 4; Table S1). We now found that each antibiotic class selected for mutations at a distinct and functionally coherent set of loci, consistent with strong parallel selection at the molecular targets of the different drugs. CIP clones in our panel recurrently albeit not uniformly harbored mutations in the DNA gyrase genes *gyrA*/*gyrB* and especially in the gene *mexS*, which is a known negative regulator of the MexEF-OprN efflux pump. Mutations in the DNA-gyrase subunits are the canonical mechanism of fluoroquinolone resistance, while MexS abrogation and resulting up-regulation of the MexEF-OprN efflux pump is a known mechanism for CIP resistance (18, 34). These same targets have been repeatedly observed in resistant isolates from patients and experimental evolution studies (35–38).

**Figure 4.**
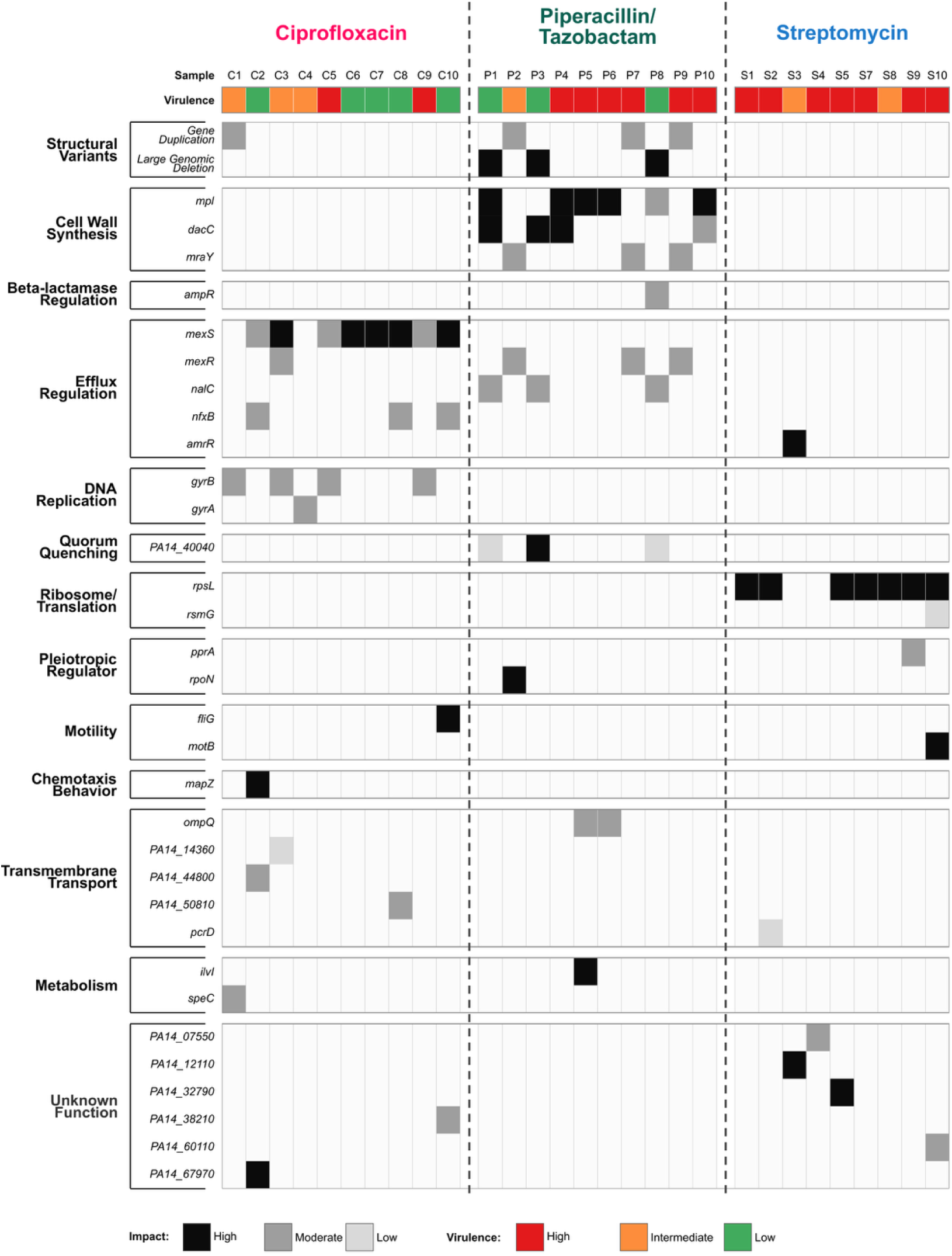
Heatmap showing different types of mutations categorized by their functions. Samples are color-coded according to their virulence phenotype observed in the acute killing assay. Black tiles indicate high-impact mutations (e.g., frameshift or stop-gained), grey moderate-impact variants (missense or small indels), and white low-impact changes (synonymous substitutions). Genes are grouped by functional category (Tables S1-S3; Figs. S4, S5) along the vertical axis. The various clones are shown along the horizontal axis.

PIT clones showed parallel mutations in three genes involved in cell wall synthesis, which is a recognized route to β-lactam resistance in *P. aeruginosa*, and has been reported as a repeatable evolutionary response to β-lactam exposure (38–40). Moreover, evolved PIT resistance is further associated with several additional known resistance genes, including *ampR*, a regulator of the ampC β-lactamase, and the efflux pump regulatory genes *nalC* and *mexR* (22, 34, 40). Most STR clones carried *rpsL* substitutions, consistent with the well-established role of ribosomal protein S12 mutations in high-level streptomycin resistance. The recurrence of *rpsL* changes across independent lineages indicates strong, predictable selection on the translation apparatus under aminoglycoside pressure (40, 41).

To complement the SNP– and indel-based mutational landscape, we further investigated larger structural variants (SVs). We identified two SVs in the sequenced clones. One was a duplication of a chromosomal region spanning 4.345–4.355 Mb of the PA14 genome, corresponding to a segment of the Pf5 prophage (Table S2). Amplification of this approximately 10-kb locus was detected in clones C1, P2, P7, P9 that were isolated from different resistance backgrounds (Fig. 4). The duplication encompassed genes associated with phage assembly, regulatory functions, and stress-response pathways (Fig. S4). Such prophage amplifications have been reported to alter stress responses, regulatory networks, and horizontal-gene dynamics, and prophage-associated genes have been implicated in stress tolerance and adaptive phenotypes in *P. aeruginosa* (42). In addition to the duplication, we also found a large chromosomal deletion present in three PIT-evolved clones (P1, P3, and P8). This deletion removed a genomic region containing known regulators of efflux pumps as well as multiple biosynthetic gene clusters (Table S3) implicated in diverse ecological and virulence-related functions of *P. aeruginosa*, including pathways involved in biofilm formation, quorum-sensing–regulated secretion systems, toxin biosynthesis, siderophore production, pyomelanin production, and stress responses (Fig. S5, Table S3).

Together, our genomic analysis indicates that AMR evolution proceeds through antibiotic-specific trajectories involving both point mutations and large-scale genome rearrangements. For each antibiotic selection environment, we observed substantial parallel genomic changes, yet also some variation that may then explain some of the observed phenotypic differences.

### Multivariate analysis links phenotypic and genomic divergence across antibiotic-evolved clones

To integrate phenotypic and genomic variation across independently evolved clones, we applied a multivariate analysis framework incorporating growth characteristics, virulence-associated traits, infection outcomes in *C. elegans*, antibiotic resistance levels, and functional mutation categories derived from whole-genome sequencing. Two types of approaches were used: Principal component analysis (PCA) shown in the main text (Fig. 5, see also Supplement Statistical Results) and principal coordinate analysis (PCoA), shown in the supplement (Fig. S6; Supplement Statistical Results). Our PCA revealed coordinated divergence across the considered biological dimensions (Fig. 5). The first two principal components explained 44% of the total variance (PC1 = 23.6%, PC2 = 20.3%). Projection of clones onto this reduced-dimensional space revealed clear structuring according to antibiotic selection regime, with CIP-, PIT-, and STR-evolved populations occupying distinct regions of the phenotypic–genomic space. Permutation-based multivariate analysis confirmed that the antibiotic environment accounted for approximately 36.5% of the overall variation (PERMANOVA, *p* = 0.001), highlighting that the antibiotic mediated selection is a major determinant of the inferred landscape.

**Figure 5.**
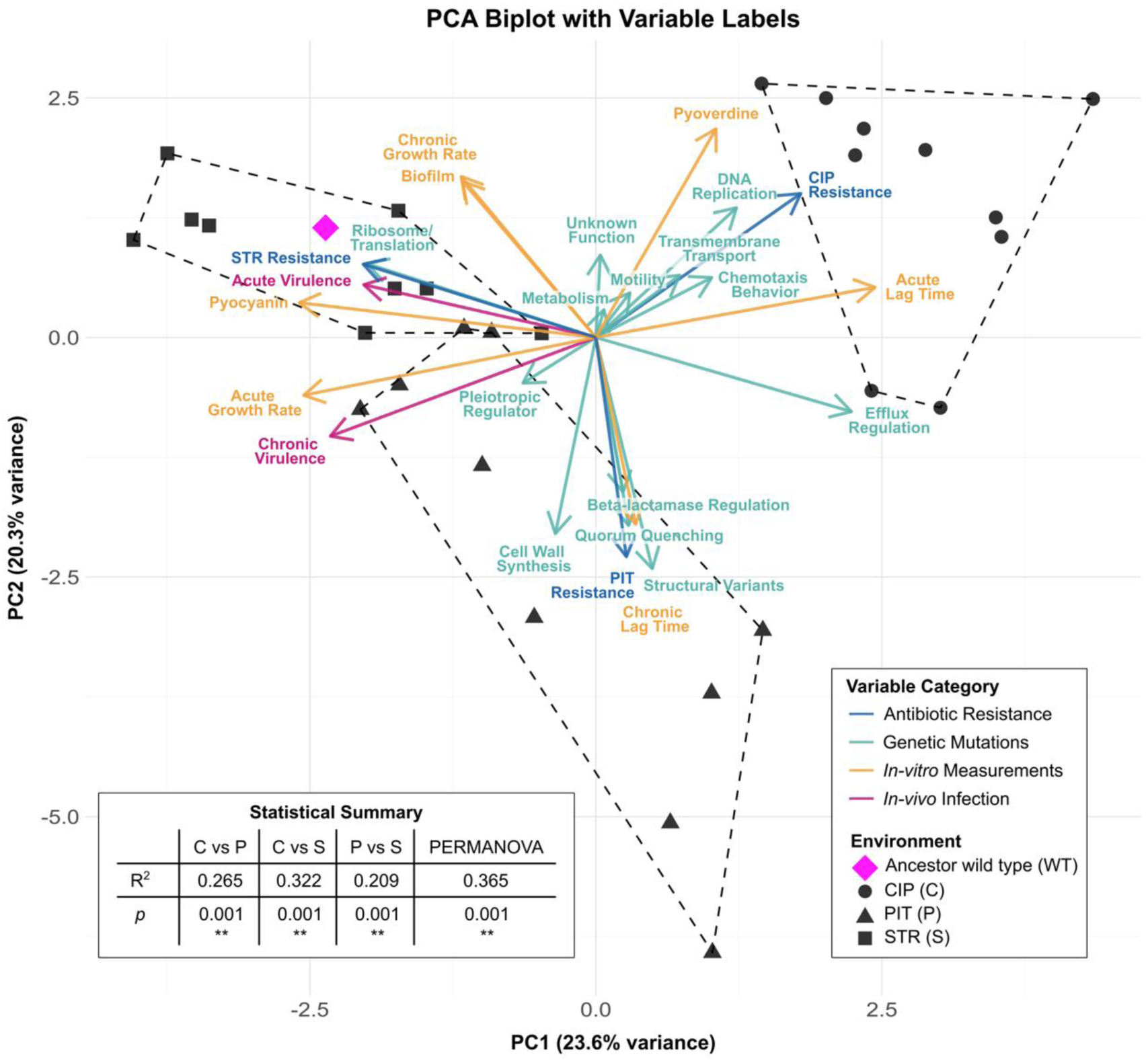
Integrated multivariate analysis of phenotypic and genomic divergence in antibiotic-evolved *P. aeruginosa* clones. A Principal Component Analysis (PCA) was used to integrate *in-vitro* phenotypic traits, *in-vivo* virulence outcomes, and *in-silico* genomic features for PA14 clones evolved under ciprofloxacin (CIP; circles), piperacillin/tazobactam (PIT; triangles), and streptomycin (STR; squares) selection. The ancestral PA14 wild-type strain is shown as a magenta diamond. The PCA is based on a combined dataset capturing growth characteristics, biofilm formation, pigment production, *C. elegans* virulence traits, and functional gene-effect scores derived from whole-genome sequencing. Arrows represent variable loadings and are colored by data category: functional gene effects (cyan), *in-vivo* virulence traits (magenta), and *in-vitro* phenotypic traits (orange). Functional gene categories were quantified using cumulative weighted scores based on SnpEff impact predictions (63), where mutations within the same functional category were summed and weighted by predicted effect severity. For each clone, the resulting score reflects the cumulative mutational burden affecting that functional category. See also details in the Supplement Statistical Results and additionally the results of the corresponding principal coordinate analysis in Fig. S6.

PC1 separated evolved resistance against CIP from that against the other two antibiotics, while PC2 differentiated resistances against either STR or PIT. Loadings of individual variables indicated that PC1 was driven largely by traits associated with bacterial growth dynamics and virulence, including acute growth rate, acute and chronic virulence in the host model, production of QS–regulated pyocyanin, and efflux regulation. In contrast, PC2 was primarily associated with chronic infection–related growth dynamics, pyoverdine production, biofilm formation, and structural variants. This integrated data analysis thereby revealed that CIP resistance evolution shows a simultaneous negative relationship with acute and chronic virulence, pyocyanin production, and growth rates in acute medium, while also being positively related to pyoverdine production, and genetic changes affecting DNA replication and efflux regulation (Fig. 5). Moreover, evolved PIT resistance is most closely linked to genetic changes in cell wall synthesis, β-lactamase regulation, structural variants, and a reduction in biofilm formation, growth rate in the chronic medium, and also pyoverdine production. Evolved STR resistance is closest to the ancestor (i.e., the magenta diamond, Fig. 5), while also being positively related to acute and chronic virulence, acute and chronic growth rate, pyocyanin production, and genetic changes affecting ribosome function. Hence, CIP and STR resistance evolution appears to lead to contrasting trait associations.

The complementary PCoA (Fig. S6) recapitulated the major clustering patterns and trait relationships observed in the PCA, confirming the robustness of the multivariate structure. Together, these patterns indicate that AMR evolution reorganizes multiple phenotypic and genomic traits in a coordinated manner that are driven by the divergent antibiotic-dependent selective constraints.

## Discussion

AMR is usually evaluated as a stand-alone trait, most likely due to standard operating procedures of clinical diagnostics. This narrow lens on AMR misses critical connections to other life history traits that result from the involvement of pleiotropic regulators in determining AMR expression (i.e., genetic constraint or genetic trade-off) and/or a re-allocation of limited energetic resources upon AMR evolution (i.e., energetic trade-off) (3–5, 8, 9). Virulence is a key pathogen life-history trait, which can mediate access to host resources, thereby reproductive rates and transmission to new hosts, and thus it has a central influence on pathogen fitness. Virulence is obviously also of high clinical relevance, as it affects host health and survival. To date, a causal understanding of the consequences of AMR evolution on virulence is limited, because it requires information on ancestors and their antibiotic-adapted descendants. Our study addresses precisely this knowledge gap by comparing experimentally evolved pathogens with their ancestor, using whole-genome sequencing, quantitative *in-vitro* phenotyping, and two types of *in-vivo* virulence assays with a *C. elegans* infection model. Our results now demonstrate that AMR evolution in *P. aeruginosa* PA14 reorganizes growth dynamics, virulence-factor production, and infection outcomes in an antibiotic-specific way. Our work additionally reveals that for each antibiotic, the underlying trajectory of resistance evolution and thus the specific type of genetic change determines the exact characteristics of the AMR-virulence trade-off. Below, we first discuss the observed trade-offs for each type of antibiotic, followed by an integrative exploration of our findings for their clinical application.

### The evolution of fluoroquinolone resistance is positively related to virulence, yet never beyond wildtype levels

The clones with evolved resistance to the fluoroquinolone ciprofloxacin, CIP, produced trade-offs with a variety of traits. These included collateral sensitivity to both PIT and STR (Fig. 1A), reduced growth rates (Fig. 1B), reduced pyocyanin productions (Fig. 2B), reduced biofilm formation in half of the clones (Fig. 2A), reduced acute virulence in almost all of the clones (Fig. 3A), and reduced chronic virulence in more than half of the strains (Fig. 3B). Such comprehensive resistance costs were not observed for evolved resistance to the other two antibiotics. Some of the CIP-related trade-offs are supported by previous studies, including CIP resistance-related collateral sensitivities towards aminoglycosides (23, 26–28) and also CIP resistance-related growth costs (5, 23, 43). Overall, the results suggest that CIP resistance evolution causes a reduction in virulence. Although virulence levels are lower than the ancestral wildtype, the exact levels of resistance are significantly positively correlated with the degree of evolved CIP resistance (Fig. 3C). This result further suggests that at least some of the mechanisms for CIP resistance are directly connected to the expression of virulence.

CIP resistance is driven by two main types of genetic changes that either affect efflux pump regulation or DNA replication. More specifically, eight out of ten CIP-evolved clones carry loss of function mutations in *mexS*, the negative regulator of the MexEF-OprN efflux pump (Fig. 4). MexS inactivation leads to constitutive overexpression of the MexEF-OprN efflux pump, which is known to export CIP but also extrudes HHQ (4-Hydroxy-2-heptylquinoline), the biosynthetic precursors of the *Pseudomonas* quinolone signal (PQS) (34, 44). The depletion of PQS was previously shown to abrogate the expression of PQS-dependent virulence factors, including for example pyocyanin, consistent with our measured suppression of pyocyanin production (Fig. 2B). A parallel observation from clinical study corroborates this trajectory: MexEF-OprN inactivating mutations are enriched among cystic fibrosis (CF) isolates relative to acute respiratory isolates, and their loss increases QS-mediated elastase and rhamnolipid expression, enhancing virulence (12). This *mexS*-mediated change in the MexEF-OprN efflux pump activity may also account for the robust collateral sensitivity of CIP clones toward STR, following the previously proposed model for *E. coli*, for which aminoglycoside susceptibility is mediated via changes in transmembrane potential, which itself is directly linked to efflux pump activity and thus export of non-aminoglycoside antibiotics like ciprofloxacin (7, 45). Interestingly, some CIP-resistant clones retained ancestral virulence levels or did not lose virulence completely, even though they all do not produce any pyocyanin and even though almost all harbor a mutation in *mexS*. These clones C1, C3, C4, C5, and C9 exclusively share mutations in the genes *gyrA* or *gyrB*, which encode the target of the CIP antibiotic. This observation may suggest that these target gene mutations are positively linked to the expression of virulence factors beyond those measured by us and/or that they may counteract the suppression of virulence induced by overexpression of the MexEF-OprN efflux pump. Together, these findings highlight that the evolutionary path to resistance directly shapes the AMR-virulence relationship, potentially leading to contrasting virulence outcomes contingent on the affected resistance mechanism.

### Complex genetic changes explain occasional loss of virulence upon β-lactam resistance evolution

Three of the evolved clones with the highest PIT-resistance levels completely lost acute virulence (clones P1, P3, and P8), indicating a possible AMR-virulence connection. These three clones are unique, because they were also the only ones that completely lost their ability to produce pyoverdine (Fig. 2C), and that gained an ability to produce pyomelanin (Fig. 2D). These three clones also showed reduced biofilm formation and pyocyanin production, but in this case not exclusively, but instead losses shown also by other clones (Fig. 2A, 2B). Moreover, these three clones are among the only four with significant collateral sensitivity towards the aminoglycoside STR (Fig. 1A). These complex phenotypic changes strongly suggest that PIT resistance evolution in these three clones was based on an exclusive mutation in a pleiotropic gene and/or some other complex genetic change. Intriguingly, three genetic alterations are unique to these three clones (Fig. 4). The first one is likely to contribute to PIT resistance and affects the gene *nalC*, a known negative regulator of the MexAB-OprM efflux pump that exports β-lactam antibiotics, such as PIT. The second includes moderate to impactful mutations in the gene *PA14_40040*, which is predicted to encode a penicillin acylase, which likely functions in quorum quenching but not AMR (46). The third unique genetic change is a large genomic deletion (Fig. S5; Table S3) that encompasses three types of genes of relevance in the context of our study: (i) Several biosynthetic gene clusters (BGC) encoding known virulence factors such as pyoverdine, biofilm-producing exopolysaccharides, the toxin hydrogen cyanide, phenazine, and other factors contributing to adhesion and biofilm formation (21, 22) (Table S3); the deletion of these BGCs likely explains the loss of acute virulence. (ii) Genes involved in tyrosine catabolism, especially the central gene *hmgA*; its loss is known to enhance pyomelanin production (47), thereby explaining the observed pyomelanin phenotype. (iii) Several AMR-related genes, including *mexX* and *mexY*, encoding central components of the MexXY-OprM efflux pump; the loss of this efflux pump is known to increase susceptibility to aminoglycosides (34, 48), thereby likely explaining the observed collateral sensitivity towards STR but not the increase in resistance towards PIT. Taken together, the complex phenotypic changes of these three clones are likely explained by the coincidental occurrence of AMR gene mutations, conferring PIT resistance, and the large deletion, likely accounting for all other observed changes. Even though these coincidental genetic changes have likely arisen independently by chance, their joint spread may have been favored by genetic hitchhiking and clonal interference. Interestingly, all three clones maintained chronic virulence, which is apparently not influenced by the virulence mechanisms encoded within the deleted region.

The remaining clones appear to have evolved PIT resistance through mutations in genes involved cell wall synthesis (Fig. 4), yet without any or only minor effects on virulence (Figs. 2 and 3). Consequently, the overall results corroborate the role of the evolutionary trajectory leading to AMR as a critical determinant of effects on other life history characteristics, in this case apparently due to the coincidental occurrence of several genetic changes and their likely spread through hitchhiking and clonal interference.

### Aminoglycoside resistance imposes low pleiotropic costs and preserves virulence

STR-evolved clones followed yet another path than those with CIP– and PIT-resistance. All of these clones evolved very high resistance levels against STR (Fig. 1), yet only two out of the nine showed significant reductions in acute virulence (Fig. 3A; clones S3 and S8), while none varied significantly from wildtype in chronic virulence (Fig. 3B). Three out of the nine clones (clones S3, S4, S8) showed significant reductions in some of the virulence-related traits, such as biofilm formation, pyocyanin and pyoverdine production (Fig. 2A-2C). These results do not indicate a specific AMR-virulence trade-off. The three clones with reductions in virulence-associated traits do not share any genomic changes. In fact, seven out of the nine clones have likely achieved STR resistance through point mutations at codons 42 and 87 in the gene *rpsL*, encoding ribosomal protein S12 and representing known targets of aminoglycoside selection (40, 41). Evolved STR resistance is further associated in one case with an impactful *amrR* gene mutation, which should result in upregulation of the MexXY-OprM efflux pump, thereby likely explaining STR resistance (34). Overall, the general lack of virulence loss in the STR clones highlights that AMR evolution does not intrinsically impose virulence costs.

### Multivariate integration reveals an intricate AMR–virulence relationship with implications for antimicrobial stewardship and treatment decisions

We integrated the collected data using both principal component and principal coordinate analyses (i.e., PCA, PCoA), consistently highlighting three distinct clusters characterized by adaptation to either of the three antibiotics (Figs. 5, S6). These analyses thereby emphasize that these antibiotics with their different mode of action define distinct selective environments that are not only connected to distinct underlying genomic changes but also to distinct changes in virulence traits and life-history characteristics. Consequently, our results reject the idea of a universally applicable AMR-virulence trade-off. Instead, we find that resistance to some antibiotics is more likely to cause a loss of virulence (applicable to the fluoroquinolone CIP), whereas resistance to other antibiotics rather favors maintenance of virulence (true for the aminoglycoside STR).

Our findings on the AMR-virulence relationship are of particular medical relevance. In detail, they challenge any framework that evaluates pathogen risk solely through resistance profiles. Instead, we identified a multivariate trait relationship that is currently absent from clinical decision-making frameworks, in spite of its potential importance. For example, a clone that is maximally resistant to CIP may simultaneously be collaterally sensitive to STR and attenuated in acute virulence, thereby yielding a combination of traits that may be clinically acceptable, which is not the case for bacteria with evolved STR resistance that maintain high virulence levels. Moreover, our findings additionally emphasize that the AMR-virulence relationship must be interrogated empirically rather than assumed, because it depends on both the antibiotic used for initial or earlier treatment as well as the exact underlying trajectory of resistance evolution.

In conclusion, this study provides, to our knowledge, the first comprehensive, multi-drug, multi-assay dissection of evolutionary trade-offs between AMR and virulence using a controlled experimental set-up. The central finding is not a universal trade-off but an antibiotic-specific evolutionary landscape, in which the direction, magnitude, and mechanistic basis of AMR-virulence interactions are determined by which cellular process is targeted by the antibiotic and which specific resistance mechanism emerges and spreads through the pathogen population.

## Materials and Methods

### Isolation of highly resistant PA14 clones from evolved populations

All bacterial strains were derived from previously experimentally evolved populations of *P. aeruginosa* PA14 described in (23). Briefly, isogenic populations (10⁵ CFU/mL) were experimentally evolved against increasing concentrations of different antibiotics: 10 each under continuous exposure to ciprofloxacin (CIP), piperacillin/tazobactam (PIT), or streptomycin (STR). The evolved populations were stored at −80 °C. For this study, frozen stocks were revived on LB agar (37 °C, 24 h). To enrich for resistance-stabilized genotypes and remove any residual susceptible cells, populations were subsequently plated on LB agar containing 2× the minimal inhibitory concentration (MIC) of the respective antibiotic. A single colony was then picked from each plate and stored again at −80 °C. This yielded 29 clones in total: 10 from CIP, 10 from PIT, and 9 from STR lineages (STR population 6 failed to recover after thawing).

Two reference strains were carried alongside evolved clones in all assays: wild-type *P. aeruginosa* PA14 as the ancestral baseline, and *P. aeruginosa* PA14 Δ*lasR*Δ*rhlR* (49) as a virulence-attenuated control, kindly provided by the lab of Bonnie Bassler (Princeton, US). The double mutant lacks both central quorum-sensing regulators, abolishing coordinated production of extracellular virulence factors including proteases, toxins, and biosurfactants. For pyoverdine analysis, we additionally included a *ΔpvdS* strain, kindly provided by the lab of Susanne Häusler (Hannover, Germany), which no longer produces pyoverdine, as a negative control.

### Quantification of resistance levels (MIC determination) and collateral response

The antibiotic susceptibility of all evolved clones was measured using the Etest® gradient diffusion assay (Liofilchem S.r.l.). Briefly, a single colony of each clone was grown in LB broth containing antibiotic. Overnight cultures were standardized to an OD_600_ of 0.1-0.15 in PBS. Standardized suspensions were plated on LB agar containing no antibiotic. Etest strips containing gradient concentrations of CIP, PIT, STR, were applied. Plates were incubated at 37 °C for 18-24 h. MIC was read at the point where the growth inhibition ellipse intersected the printed scale on the strip.

### Growth measurements in chronic and acute killing media

Bacterial growth was measured directly in the media used for *C. elegans* infection experiments (described below) and thus under conditions used for virulence inferences. Cultures were inoculated at OD_600_ = 0.1 and incubated at 37 °C with shaking. OD_600_ was recorded every 15 min for 24 h using a plate-reader spectrophotometer (either Tecan Infinite 200 Pro, Tecan Austria GmbH; or BioTek Epoch 2, Biotek Instruments Inc./Agilent Technologies Inc.). Growth curves were fitted to the logistic growth model, using the R package Growthcurver and minpack.lm (50, 51). From fitted parameters, four summary statistics were extracted: maximum growth rate, lag time, and carrying capacity, and overall growth performance, of which the latter was quantified as the area under the curve (AUC), computed numerically using the trapezoidal rule over the full 24 h window with the R package pracma (52).

### Biofilm formation

Biofilm formation was quantified using a standard crystal violet microtiter plate assay (53). Overnight cultures were adjusted to OD_600_ = 0.1 in fresh LB, then diluted 1:100 into LB-filled wells of a 96-well plate (198 µL LB + 2 µL culture per well). Plates were incubated statically at 37 °C for 30 h. Following incubation, the planktonic phase was aspirated and each well was washed three times with sterile deionized water. Adherent biofilm was stained with 0.4% crystal violet for 20 min, washed to remove excess dye, and the retained stain was dissolved in 99% ethanol. Biofilm biomass was quantified by measuring absorbance at OD_595_.

### Quantification of pyocyanin, pyoverdine, and pyomelanin

Pyocyanin, pyoverdine, and pyomelanin are secreted *P. aeruginosa* metabolites that respectively act as a redox-active phenazine toxin under quorum-sensing control, a high-affinity fluorescent siderophore mediating iron scavenging, and a melanin-like pigment involved oxidative-stress protection and persistence. Each of these metabolites has been implicated in bacterial virulence and shown to influence survival in whole-animal models including *C. elegans* (33, 54, 55). For each clone, culture supernatants were collected from standardized overnight cultures, and pigment absorbance was measured using UV–Vis spectrophotometry at wavelengths specific to each metabolite, including for (i) pyocyanin: 695 nm (56); (ii) pyoverdine: 400 nm (57); and (iii) pyomelanin: at 470 nm, the latter corresponding to the empirically determined absorbance maximum identified by full UV-Vis spectral scanning (200–700 nm) of representative samples prior to measurement. All toxin measurements were normalized to OD_600_ to account for differences in biomass and growth rate across clones. *The ΔlasRΔrhlR* strain served as a QS-deficient negative control for pyocyanin and pyomelanin production, the *ΔpvdS* strain as a negative control for pyoverdine production.

### *In-vivo* virulence measurements

Virulence of evolved clones was assessed using *C. elegans* as an infection host, leveraging its conserved innate immune pathways and well-established utility as a bacterial pathogenicity model (24). Two mechanistically distinct assays were used: acute killing, which captures rapid toxin-mediated mortality, and chronic killing, which reflects intestinal colonization and sustained host–pathogen interaction over several days. The killing assay protocols were adapted from (24) with small modifications.

For acute killing, overnight cultures were used to spot 8 µL lawns onto peptone glucose sorbitol (PGS) agar plates (10 g Bacto proteose peptone, 27.4 g D-sorbitol, 10 g NaCl, and 50 mL of 20% D-glucose, 17 g Bacto agar in 950 mL deionized water). PGS medium induces *P. aeruginosa* toxin production and does not require intestinal colonization or even live bacteria to cause worm mortality. Plates were incubated at 37 °C for 24 h, then shifted to 25 °C for a further 24 h to induce toxin production. Twenty age-matched L4-stage N2 hermaphrodite worms were transferred onto each plate and survival was scored at 6 h post-exposure by assessing motility and response to gentle touch, with percentage mortality used as the virulence readout.

Since evolved clones varied in growth rate and since QS-regulated virulence factors in *P. aeruginosa* are cell-density–dependent, we additionally tested whether inoculum density influenced killing outcomes. Two inoculum densities were compared: OD_600_ = 0.1 (low) and OD_600_ = 1.0 (high). To assess the resulting cell numbers, bacterial lawns were scraped from PGS plates after each assay, resuspended in sterile PBS, serially diluted, and plated for CFU scoring.

For chronic killing, lawns were prepared on modified NGM agar (3.5 g Bacto proteose peptone, 3 g NaCl, 20 g Bacto agar in 1,000 mL deionized water, followed by autoclaving and subsequent addition of sterilized standard NGM supplements), yielding a low-osmolarity enriched peptone medium that supports *P. aeruginosa* intestinal colonization and slow progressive killing. Prepared plates were incubated for 48 h to allow stable lawn establishment. Before adding worms, 80 µL of PEG 2000 solution was dispensed along the lawn perimeter to prevent worms from escaping the bacterial surface (58). Thirty synchronized L4-stage worms were transferred onto each plate and survival was scored every 24 h for four consecutive days. Worms were moved daily to freshly prepared PEG-coated plates carrying the same bacterial lawn. The assay endpoint was defined as complete mortality on wild-type PA14 plates. Survival data were analyzed using Kaplan–Meier curves. To enable direct quantitative comparison across clones, survival outcomes were additionally converted to a Virulence Index (VI = 1 − [AUC_observed / AUC_maximum]), where AUC is the area under the Kaplan–Meier survival curve and AUC_maximum represents the theoretical maximum assuming complete survival across all scored time points. This approach compresses multi-day survival data into a single normalized metric that is directly comparable across clones and replicates.

### Whole genome sequence analysis

We characterized whole genome sequences of the evolved clones in comparison to the ancestor, to assess the genomic changes underlying AMR and possibly virulence. Genomic DNA was extracted from each *P. aeruginosa* clone using the Macherey-Nagel Bacterial Genomic DNA Extraction Kit (Macherey-Nagel; Düren, Germany), following the manufacturer’s guidelines. Whole-genome sequencing (150 bp paired-end) was performed by Novogene LTK, Cambridge, UK using an Illumina NovaSeq X Plus 25B platform. Sequencing data were processed using a standard variant analysis pipeline. Raw reads were quality filtered and adapter trimmed with Trimmomatic (v0.39) (59), then aligned to the *P. aeruginosa* PA14 reference genome using Bowtie2 (v2.5.1) (60). BAM files were processed with SAMtools (v1.17) (61) and Picard (Broad Institute toolkit) to format alignments, mark duplicates, and add read group information. SNPs and small INDELs were identified with bcftools (v1.14), followed by filtering based on quality and allele frequency, and comparison to the ancestral PA14 strain. Genome-wide coverage depth and uniformity were evaluated using deepTools (62). Structural variants were detected using the Structural Variation Engine (SVE), incorporating SNVnator (v0.3.3), DELLY (v2), and LUMPY, with consensus calls retained using SURVIVOR. Variants present in the reference genome background were excluded. Finally, variants were annotated with SnpEff (63) to predict their impact on coding sequences and regulatory elements, including genes associated with AMR and virulence pathways.

## Statistical analyses

All statistical analyses were performed in R (v4.x) (64). Phenotypic measurements (e.g., virulence outcomes, growth parameters, biofilm biomass, and toxin production) were compared against the wild-type PA14 ancestor using Dunnett’s multiple comparison test, implemented via the multcomp package (65), to control family-wise error rate across evolved clone groups. To summarize and visualize multivariate phenotypic diversity across antibiotic selection backgrounds, we applied both principal component analysis (PCA) and principal coordinates analysis (PCoA). PCA was used for continuous variables, integrating *in-vitro* traits (growth kinetics, MIC profiles, biofilm formation, pyocyanin production), *in-vivo* virulence outcomes from both acute and chronic killing assays, and functional genomic features derived from variant annotation. Rather than including raw mutation counts, SNPs and structural variants were grouped into biologically meaningful categories (e.g., cell wall synthesis, efflux regulation, DNA replication, etc.), and the predicted impact scores of all mutations within each category were summed per clone, yielding continuous functional burden values suitable for PCA. PCoA was applied separately to binary-encoded genomic data, where each functional category was scored as 1 (at least one mutation detected) or 0 (no mutation detected), allowing multivariate patterns to be summarized based on the distribution of functional mutation categories independent of mutation severity. To formally test whether clones evolved under different antibiotics occupied statistically distinct multivariate spaces, PERMANOVA was performed using the adonis function in the vegan package, assessing both overall group separation (CIP vs. PIT vs. STR) and all pairwise contrasts (CIP vs. PIT, PIT vs. STR, STR vs. CIP).

## Data availability

The source data of this study is provided in a supplementary Source Data file. The statistical results are shown in a Supplementary Statistical Results file. Summaries of the results of the whole genome sequencing analysis are provided as Tables S1-S3. The Illumina sequencing reads have been submitted to the Sequence Read Archive under BioProject PRJNA1345347.

## Supporting information

Supplement Statistical Results

## Acknowledgments

We thank Prof. Bonnie L. Bassler (Princeton University, US) for providing the *P. aeruginosa* PA14 ΔlasRΔrhlR double mutant strain, Prof. Susanne Häußler (Helmholtz Centre for Infection Research, Hannover, Germany) for providing the ΔpvdS mutant, and the Schulenburg group for helpful discussions and feedback.

## Author Contributions

Conceptualization: SP, BP, HS; formal analysis: SP, MH, BP, HS; funding acquisition: HS; investigation and methodology: SP, AM, MH, BP; supervision: BP, HS; writing of original draft: SP, BP, HS; review and editing: all authors.

## Classification

Biological sciences, Evolution

