## Supplementary Figures S1-S6 for "Evolutionary trade-offs between antimicrobial resistance and virulence in *Pseudomonas aeruginosa*"

**This PDF file includes:**

Supplementary Figures S1-S6

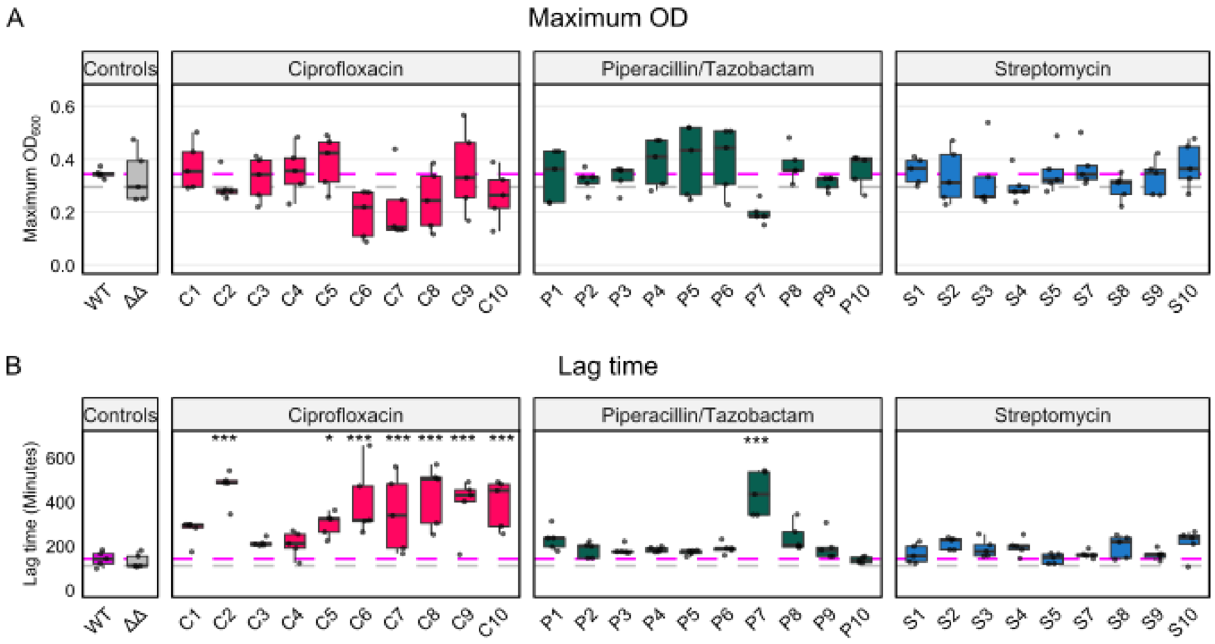


**Supplementary Figure S1. Growth performance of evolved *P. aeruginosa* PA14 clones in drug-free medium relevant to the acute-killing environment.** Growth characteristics were quantified in the same medium used for the *C. elegans* acute-killing assay (n = 5), using growth dynamics as a proxy for bacterial fitness. (A) Maximum carrying capacity measured as maximum OD (K). (B) Lag phase duration in minutes). The magenta dashed line denotes the median value of the wild-type PA14 strain (WT). The grey dashed line denotes the median of the Δ*lasR* Δ*rhlR* (ΔΔ) mutant. Statistical comparisons were performed using Dunnett's test against the WT. Significance thresholds are indicated as *$p < 0.05$, **$p < 0.01$, and ***$p < 0.001$ (see also Supplement Statistical Results).

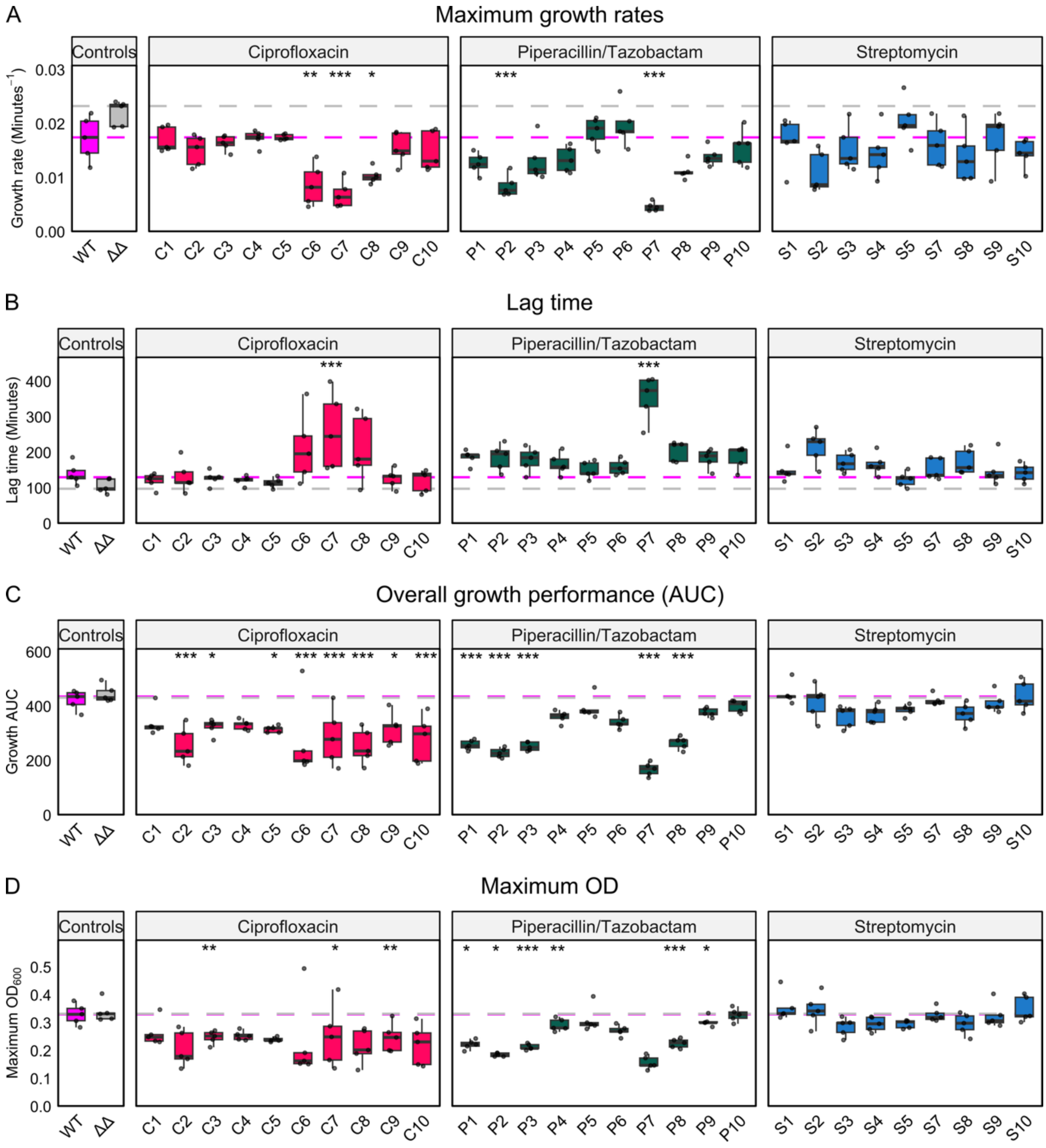


**Supplementary Figure S2. Growth performance of evolved *P. aeruginosa* PA14 clones in drug-free medium relevant to the chronic-killing environment.** Fitness landscapes of all evolved clones were quantified in the same growth medium used for the *C. elegans* chronic-killing assay (n = 5). Four quantitative metrics were extracted: (A) maximum growth rate (μ), (B) lag phase duration, (C) cumulative growth (AUC), and (D) maximum carrying capacity (K). The magenta dashed line denotes the median value of the wild-type PA14 strain (WT). The grey dashed line denotes the median of the Δ*lasR* Δ*rhlR* (ΔΔ) mutant. Statistical comparisons were performed using Dunnett's test against WT (**p* < 0.05; ***p* < 0.01; ****p* < 0.001; see Supplement Statistical Results). In contrast to acute-killing, several CIP-evolved clones — specifically C1, C2, C3, C4, C5, C9, and C10 — displayed WT-like growth rates accompanied by reduced lag times.

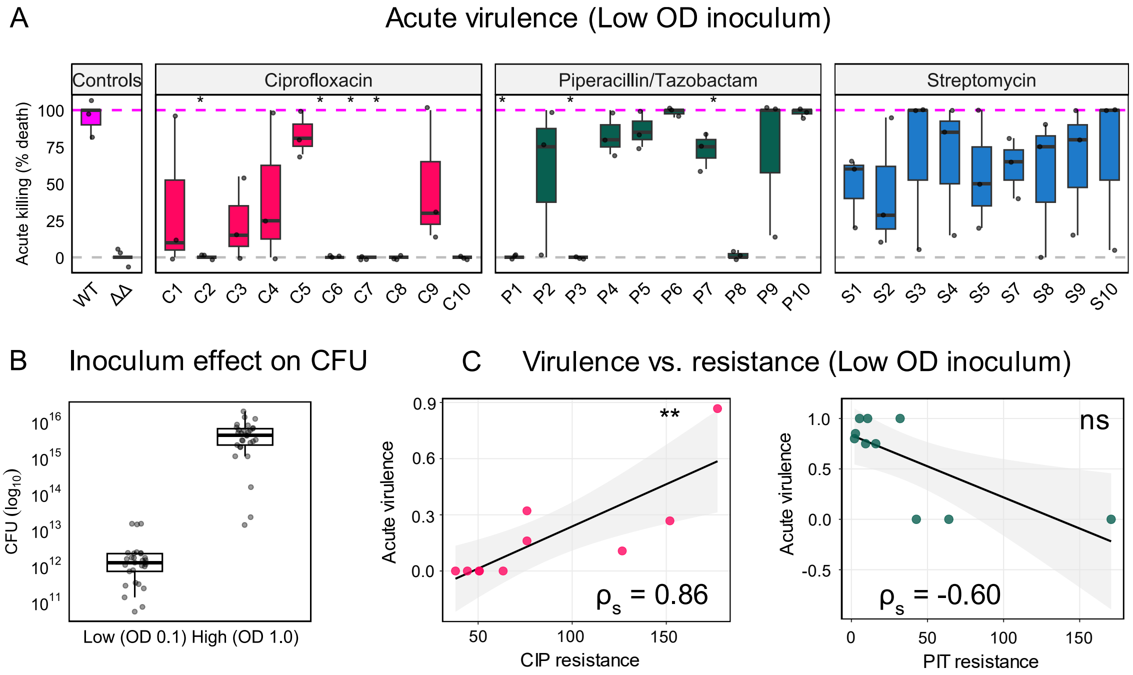


**Supplementary Figure S3. Effect of low inoculum on acute-killing outcomes.** (A) Acute-killing assay outcomes using low initial inoculum (culture diluted to $OD_{600}$ = 0.1 prior to plating). Virulence is shown as the percentage of worms dead after 6 h post infection. The magenta dashed line denotes the median value of the wild-type PA14 strain (WT). The grey dashed line denotes the median of the Δ*lasR* Δ*rhlR* (ΔΔ) mutant. Statistical comparisons were performed using Dunnett's test against WT (**p* < 0.05; ***p* < 0.01; ****p* < 0.001; see Supplement Statistical Results). Across both inoculum regimes, high (Fig. 3A) and low OD, clones displayed comparable virulence patterns, indicating that initial inoculum size is not a determining factor for acute-killing outcomes. (B) Validation of inoculum preparations. CFU quantification was performed directly from acute-killing assay plates prepared using high- and low-inoculum regimes. Note that the y-axis is $log_{10}$ transformed. Low inoculum preparations consistently yielded lower CFU densities, confirming that the two conditions establish reproducibly distinct bacterial loads. (C) Correlation between antibiotic resistance level (MIC fold-change relative to PA14 WT) and virulence at low inocula in acute infection. CIP-evolved clones exhibited a significant positive correlation between resistance level and virulence under both inoculum conditions (Spearman rank correlation; n = 10). PIT-evolved clones displayed a significant negative correlation between resistance and virulence under both inoculum conditions (n = 10), consistent with a resistance–virulence trade-off.

Structural variants: prophage gene duplication in C1, P2, P7, and P9

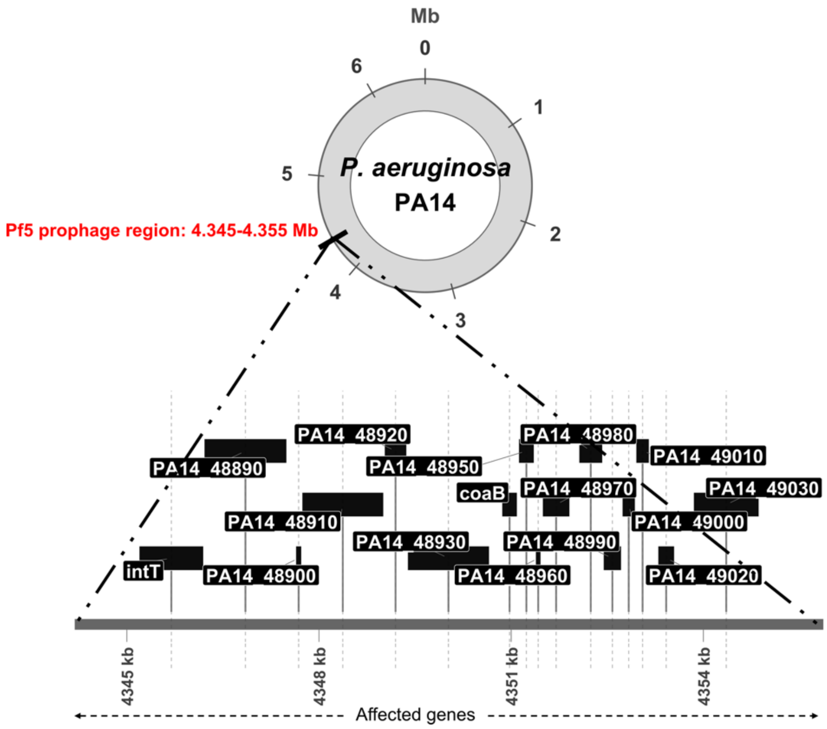


**Supplementary Figure S4. Gene duplication events in some PA14 resistant clones.** Circular representation of the PA14 genome highlighting the duplicated region (4.345–4.355 Mb; shown in black). This segment corresponds to the Pf5 prophage locus, which is amplified in clones C1, P2, P7, and P9. Expanded view of the duplicated region showing the gene content within the amplified Pf5 prophage cluster.

Structural variants: large genomic deletion in P1, P3, and P8

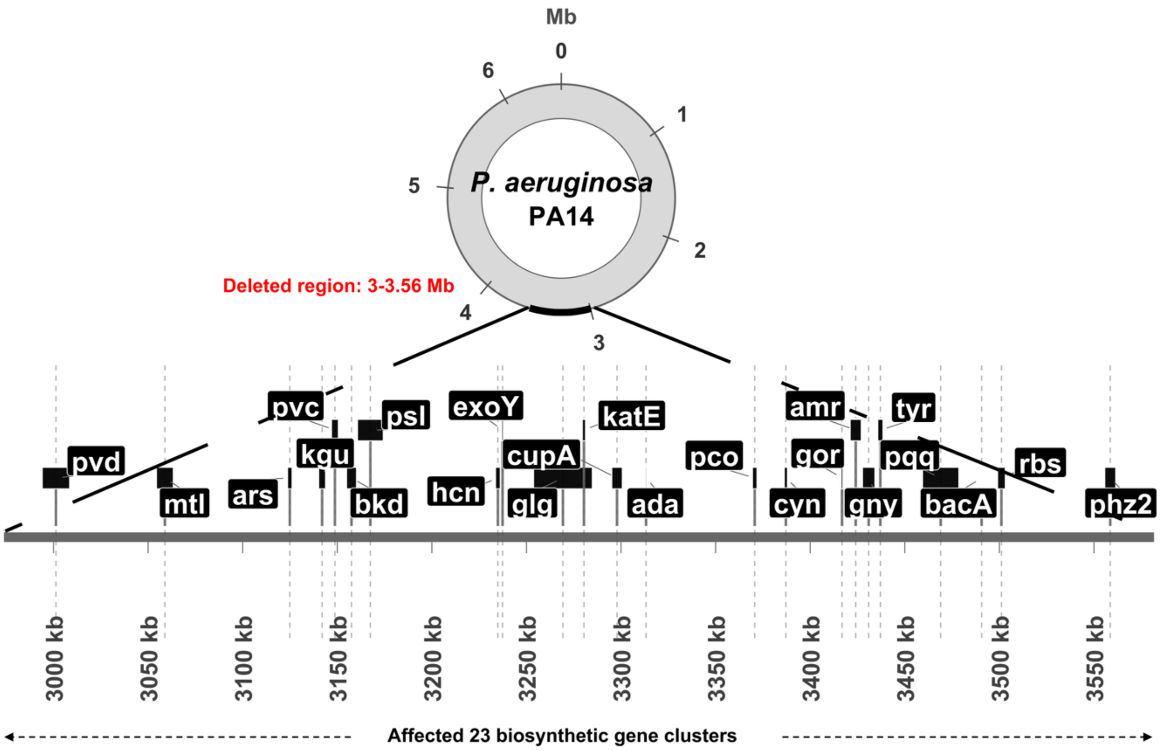


**Supplementary Figure S5**. **Large genomic deletions identified in PIT-evolved PA14 clones.** Circular PA14 genome map highlighting the deleted region of approx. 560 kb present in clones P1, P3, and P8. The deleted segment impacted a large multi-gene locus encompassing 23 biosynthetic clusters. Zoomed-in view of the deleted genomic region, illustrating several of the affected biosynthetic clusters.

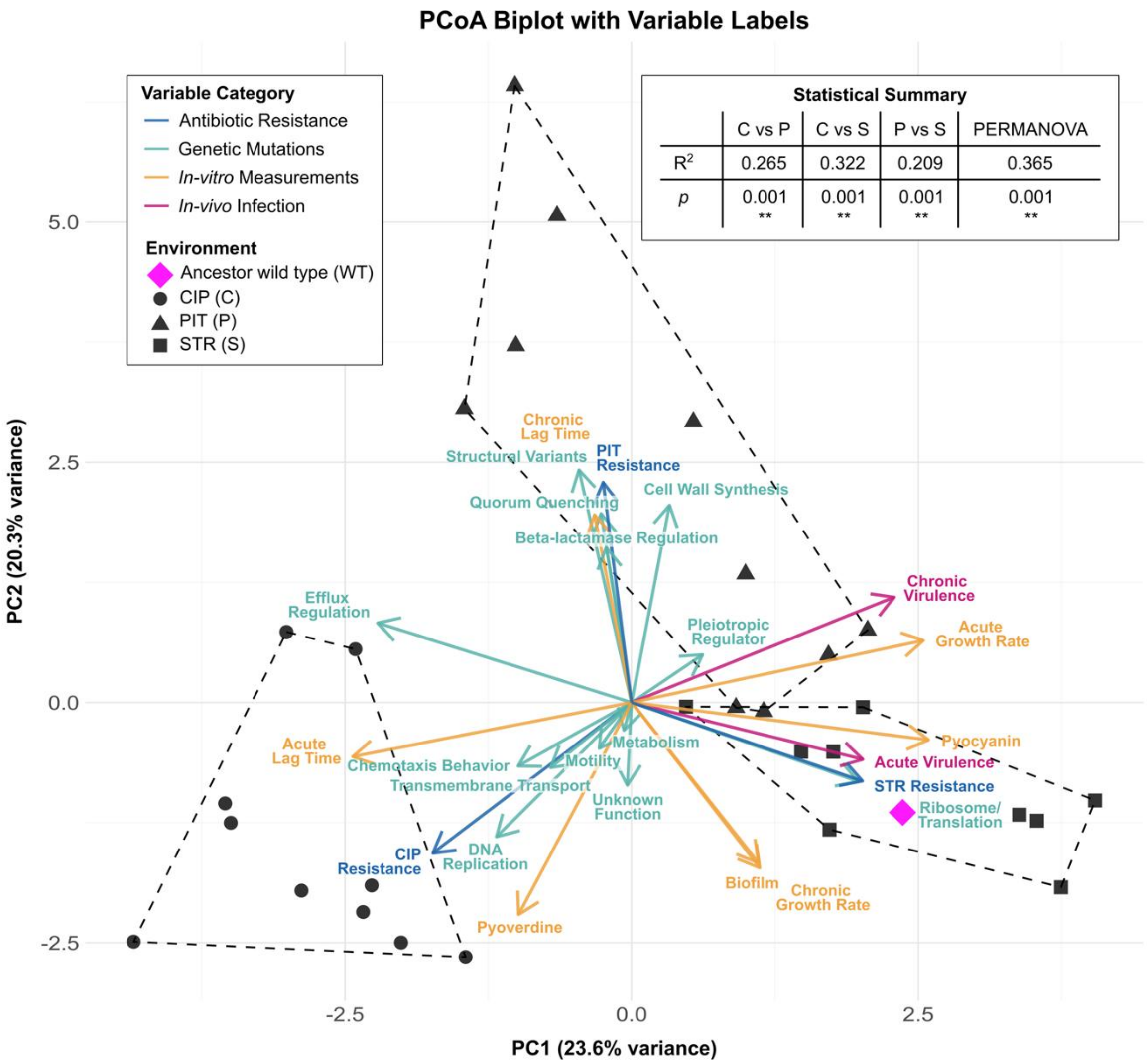


**Supplementary Figure S6: Multivariate integration of phenotypic traits, virulence outcomes, and genomic features.** Principal coordinate analysis (PCoA) integrating *in-vitro* phenotypic traits, *in-vivo* virulence outcomes, and *in-silico* genomic features for PA14 clones evolved under ciprofloxacin (CIP; circles), piperacillin/tazobactam (PIT; triangles), and streptomycin (STR; squares) selection. The ancestral PA14 wild-type strain is shown as a magenta diamond. Variable arrows are colored by measurement category. Functional gene effects were encoded using a binary scoring scheme, where a value of 1 indicates the presence of any mutation affecting a given functional category (e. g., DNA repair, efflux, motility), and 0 indicates absence of mutations in that category. The inset (top right) shows results from a PERMANOVA analysis testing the effect of antibiotic selection regime (CIP, PIT, STR, WT) on the integrated multivariate dataset. The reported $R^2$ value indicates the proportion of total variance explained by antibiotic background (****p* < 0.001 based on permutation testing), confirming that the selection environment is a major driver of coordinated phenotypic and genomic divergence (see also Supplement Statistical Results).
